# A discontinuously coupled network of phase oscillators replicate actomyosin cooperation

**DOI:** 10.1101/2023.12.04.569886

**Authors:** Benjamin Warmington, Jonathan Rossiter, Hermes Bloomfield-Gadêlha

## Abstract

Groups of non-processive myosin motors exhibit complex and non-linear behaviors when binding to actin. These operate at larger scales and time frames than an individual motor, indicating the presence of a strong cooperative disposition. Limits in contemporary microscopy prevent verification of motor-filament binding dynamics, whilst mathematical models rely on continuum abstractions in which cooperativity is implicit and individual motor behavior cannot be separated from the behaviour of the whole. Understanding the fundamental interactions driving the emergent behaviour in actomyosin therefore remains an open question. Here we suggest that the diversity of empirically observed *in-vitro* oscillations can be explained by a minimal Kuramoto-style phase oscillator model of actomyosin, where cooperativity is orchestrated by the actomyosin geometry and mechanical environment. The model mirrors the irregular and regular saw-tooth oscillations present in *in-vitro* actomyosin and sarcomeric ‘SPOC’ experiments with only adjustments of the external mechanical environment, and despite the model’s simplicity. Actomyosin-like behaviour thus arises as a generic property of the discontinuous mechanical coupling in an incommensurate architecture, rather than specific to molecular motor reaction kinetics. We demonstrate a range of synchronising behaviours arising from the cooperative motor dynamics that, once synchronised, are stable over a large range of external forces. These synchronising behaviours arise from the cooperative motor dynamics that, once synchronised, are stable over a large range of external forces. The nature of the synchronisation patterns allow recruitment of rotors as the external force increases, reducing variance in the backbone’s velocity. This is a demonstration of morphological control. Due to interest in this behaviour in contemporary robotics, we build a physical experiment, using electric motors to power our oscillators. Using the experiment we verify both the organisational and control properties of the system. This demonstrates non-biological motors can cooperate similarly to biological motors when working within an actomyosin geometry, suggesting that the actomyosin complex may not depend on motor-specific qualities to achieve its biological function. These findings offer novel insights into synchronising networks of oscillators and have potential applications in emulating actomyosin-like behaviors within contemporary robotics using non-biological motors.

## I. INTRODUCTION

Actomyosin systems are ubiquitous in eukaryotic cells, where they are responsible for a range of roles from molecule transport [1] to cell mechano-transduction [2] and cell division [3]. The active element of the system, myosin, is a bipolar active unit that binds to actin and undergoes a conformational change with the addition of ATP to march along the periodic binding sites of the actin filament [4]. One of the most studied roles of actomyosin is in muscle cells, where they provide the essential function of animal movement [4, 5]. Muscle based myosin is non-processive, where myosin cofilaments - thick bundles of active myosin molecules - form a hexagonally arranged lattice that overlaps a similar lattice of actin [6]. In the presence of ATP and excess calcium ions (analogous to turning a muscle cell on) [7] binding sites on the actin filament are uncovered, allowing the myosin heads to pull the two lattices together and contract the muscle.

The myosin system therefore predominantly has the control of activation level only via concentration of calcium and ATP, and yet the system is capable of different behaviours - regular [8] or irregular [9] triangular oscillations, static isometric tension [10], rectangular oscillations [11] - depending on the stiffness of the substrate it is acting in and the external forces being applied to it. Alongside this, independent myosin heads must coordinate with each other to allow a consistent and predictable strain rate [12–14]. This self-adaptive and organising behaviour suggests at a molecular level there are rich coupling interactions occurring between the myosin heads.

Current microscopy techniques do not allow direct visualisation of systems of individual myosin heads during contraction, preventing an empirical based understanding on the nature the different motor coupling that may be involved. Despite this limitation, *in vitro* experiments have been able to display single myosin molecule behaviours [15, 16] and the displacement behaviours of an actin filament within an actomyosin system [9, 12]. It is therefore these experiments current models aim to replicate and are measured against, initially using the two-state model of actomyosin attachment first proposed by Julicher and Prost in 1995 [17], and later using iterant forms of this model [5, 9, 18, 19]. The two-state model and its derivatives are continuous models, describing rates of change of populations of attached myosin dictated by a prescribed sawtooth binding potential. Subsequently these models provide little illumination into the emergent myosin dynamics from individual to collective motor action in the context of a contracting actomyosin system, with self-oscillations prescribed implicitly [9, 17–19]. Furthermore they cannot illuminate whether the geometrical design and spacial structure of actomyosin system play any role on the syncronization phenomenon of the motors. In 2017 Kaya *et al* [12] proposed a more complex, discrete model of myosin wherein each myosin head cycled through six separate states. This replicates to a high degree of accuracy the cooperative action of 17 interacting myosin heads in the specific experiment it models, however it does not take into account the periodicity of actin binding sites, and, as such, any effects geometry that may be incurred on the cooperative dynamics of motors, including variations in the mechanical environment or in the number of interacting myosin heads.

A separate line of enquiry using a dynamical systems approach with coupled amplitude oscillators offers numerous opportunities for modelling individual to collective molecular motor dynamics, as have been successfully exploited in a multitude of biological synchronization phenomena [20–24]. This has not yet been explored in the context of actomyosin oscillations. Kuramoto systems allow direct interrogation on how coupling regulates distinct synchronization patterns, without delving into the complicated dynamics that govern each phase oscillator, of which myosin has many. Altogether, this places an adapted form of Kuramoto’s synchronization model as an ideal mathematical framework to tackle synchronization in actomyosin systems. Here, we thus bridge this gap and investigate whether the mechanical coupling provided by the actomyosin complex, rather than any form of complex interactions arising from the molecular motor reaction kinetics, is sufficient to generate typical actomyosin oscillations observed empirically (Fig. 2). This approach also allows us to fill a gap in contemporary research in predicting how the nanometer scale dynamics of a myosin head leads to the micron scale emergent behaviours of large number of interacting motors in actomyosin systems.

The actomyosin synchronization model presented here is founded on Kuramoto’s phase oscillator model, but fundamentally differs from classical Kuramoto systems. Classically, coupling among oscillatory nodes is present at all times and dictated by a homogeneous and dense connectivity of the network, where nearby neighbours are always connected [25]. Actomyosin complexes on the other hand only couple myosin heads (our nodes) for brief periods of time, when myosin is bound and forcing the actin filament. Alongside this, any oscillatory node on the actomyosin network can be directly connected to each other regardless of how distant these nodes might be within the actomyosin complex (Fig.1 d. The actomyosin structure orchestrates the coupling dynamics and synchronization among nodes, either by linking or disconnecting nodes intermittently. Due to this discontinuous nature in coupling, the actomyosin complex is more akin to time-dependent networks [26, 27]. Our model proposes a simple, discrete model of actomyosin, where a discontinuously coupled network of mechanical phase oscillators (also denoted simply as rotors) replace myosin as an active motor unit (Fig. 1 c), and myosin’s conformational changes are represented in a continuous, cyclical manner (Fig. 1 c). Actin binding sites are modelled as equidistantly rigidly connected ‘teeth’ which are constrained to one dimensional movement and experience a resistive friction force within the actomyosin structure (Fig. 1 d). Following the suggestion by Brizandine *et al*. [28] that actin velocity is not limited by detachment rate we can expect no negative force generation from myosins. We therefore suggest simple contact forces rather than any form of more complex binding kinetics, are sufficient to generate typical actomyosin oscillations observed empirically. As such motor binding occurring via direct contact forces between the rotors and teeth (Fig. 1 b) is a key aspect of this model. The motors are only coupled for short periods of time via self-organized teeth-rotor contact mechanics through the backbone (Fig. 1 d). Another fundamental distinction of our model, also when compared with canonical Kuramoto phase oscillators [20, 21], is the fact that the rotors here are mechanical rotor entities (Fig. 1 b-d) and, as such, follow a general force-velocity relationship of molecular motors, typically characterised by linear decay (Fig. 1 a) [12, 29]. This provides the actomyosin system the potential for cooperativity via momentum balance (see Methods). The actomyosin rotor dynamical system is thus characterised by a discontinuous, mechanical coupling of individual phase oscillators embedded within the specific actomyosin architecture (Fig. 1 d). As such, we have named this the discontinuously coupled rotor (DCR) model of myosin cooperation.

**FIG. 1.**
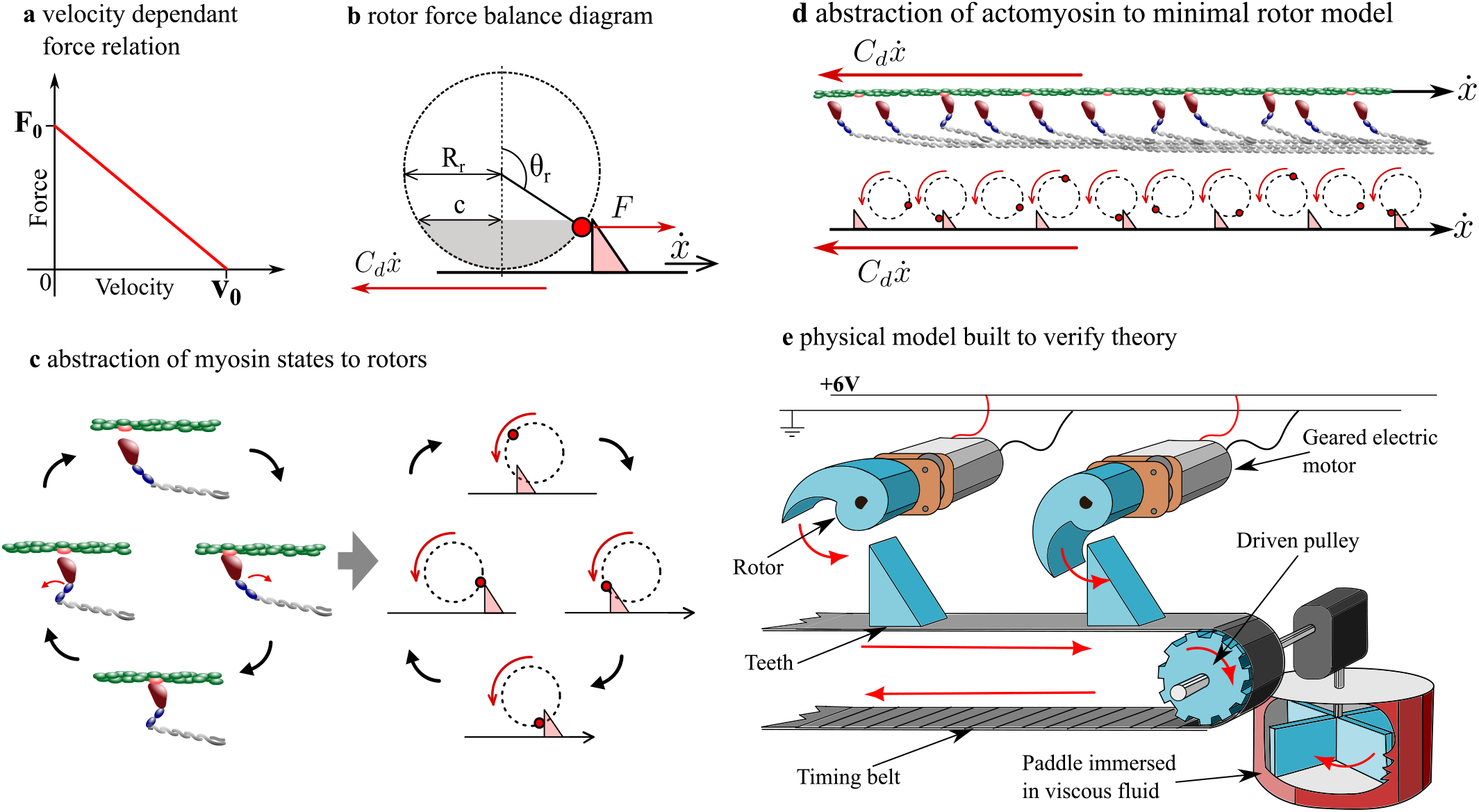
**a** We use a simple linear decay description of the molecular motor force velocity relation. **b** A free body diagram of a rotor and tooth, showing relevant parameters. The shaded area is the part of the orbit the rotor can physically come into contact with the tooth. **c** The discrete phases of myosin binding are abstracted as different parts of a rotors’ orbit. **d** A full interacting actomyosin filament is abstracted as a strip of teeth that can move through one dimension being pushed by rotors whose centre of rotation is fixed in the lab frame. The rotors are acting against a viscous drag force. **e** An experimental representation of the model was constructed as a proof of concept. A row of periodically spaced motors drive rotors that interact with teeth fixed to a timing belt. The timing belt drives a pulley that in turn drives a paddle immersed in a high viscosity fluid.

In proposing the myosin DCR model we are making several hypotheses - firstly that despite the short episodes of contact the system can provide enough mechanical feedback to entrain. We show this to be true. We show a rich umbrella of cooperative behaviours by building a simulation within observed parameters in actomyosin experiments [9, 12, 30–32]. We go on to hypothesise the co- operating behaviours of actomyosin systems, particularly regular and irregular saw tooth oscillations (Fig. 2 a,b), can be captured by any system that has regularly spaced force producing units acting on and coupled through a backbone with incommensurate regularly spaced binding sites. Thus, we show this is a scale invariant, generic property of actomyosin-style discontinuous mechanical coupling rather than specific to a molecular motor constitutive relations. As such, actomyosin-style motor cooperation can be achieved in engineering and artificial systems without requiring biological molecular motors and their intricate reaction kinetics, which may also appeal to the field of robotics.

**FIG. 2.**
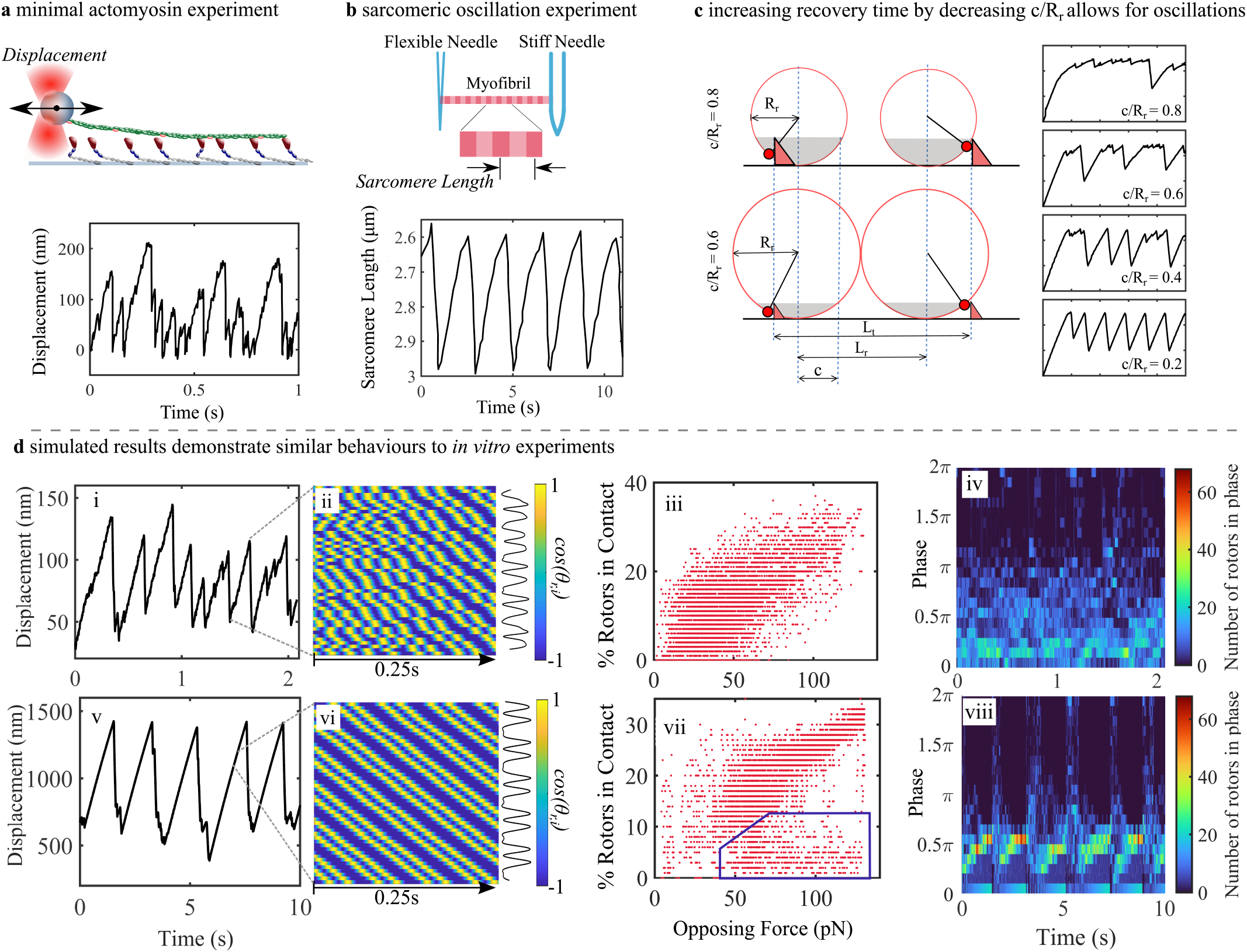
**a** diagram of minimal actomyosin experiment originally described by *Placais et al* [9]. A micron diameter bead is bonded to an actin filament and suspended in a laser trap, which acts as an elastic returning force. Myosin acts on actin to spontaneously oscillate the bead. Displacement profile taken from [12]. **b** Sarcomeres display regular oscillations at partial activation under isotonic and auxotonic conditions, as first shown by *Fabiato and Fabiato* [33]. The experimental setup and data here is taken from *Kono et al’s* recent experiments [34]. **c** We can increase the non-contactable time of a motor by increasing the size of the rotor’s radius while keeping the dimensional size of the contactable region the same. As the length scales are non-dimensionalised by the radius all the values need to be appropriately recalculated for the dimensionless model. Increasing the length of the non-contactable region mimics the relatively small time an individual myosin spends bound to actin in actomyosin, and results in more reliable oscillations. **d, i** Displacement graph for *α* = 1.2, *β* = 3.8 chosen as qualitatively similar to minimal actomyosin system (**a**). **d, ii-iv** Kymograph of the cosine of the rotors’ angle against vertical over a 0.25*s* period, scatter-graph of percentage of rotors in contact against the elastic opposing force on the filament for each time step of the whole signal segment, probability density for the neighboring rotor phase differences throughout the segment. **d, v** Displacement graph for *α* = 2.4, *β* = 0.5 chosen as qualitatively similar to sarcomeric system (**b**). **d, vi-viii** The same graphs as **d, ii-iv**, but for the signal segment in **d, v**.

We show that the proposed DCR model qualitatively replicates actomyosin behaviours, including the quintessential nonlinear, irregular saw tooth oscillations seen in minimal actomyosin systems [9, 12] and more regular sarcomeric SPOCs [8, 33, 34]. These cooperative behaviours could be induced by simply tuning the elastic forces acting against the motors and keeping the same number of motors, whilst the excellent comparisons with experiments indicate the validity of the DCR model and the assumptions invoked. Crucially, having access to the individual motor dynamics, we also show that self-organisation of individual motors lead to a diverse range of metachronal travelling waves along the actomyosin structure, bridging nano-to-micron behaviours. We demonstrate that synchronised motor systems are capable of generating much greater force, as well as minimising velocity changes from varying force by engaging or disengaging the number of motors in contact, akin to an auto control mechanism.

Finally, following our claim that this particular mechanical system has some application within the field of robotics we construct a physical experiment showing some of the key behaviours present in the simulations (Fig.1 e). Key within the behaviours demonstrated are the organisation for the motors into metachronal travelling waves and the process by which motors are engaged or disengaged as the resistive force is increased or decreased.

## II. RESULTS

In this model myosin II molecules are simplified to minimal discontinuous Kuramoto rotor equivalents (Fig. 1 c), where each rotor moves through a ‘contactable’ region and a ‘non-contactable’ region (see methods). Actin is simplified to a periodic series of ‘teeth’, which can be pushed by a rotor if the rotor meets the tooth within the contactable region (Fig. 1 b). Physiological style binding is ignored in favour of contact forces between the rotor and the tooth - as the rotors only move in an anticlock-wise direction we say that once a rotor contacts a tooth it is ‘engaged’ as it will release only once the tooth moves outside of a rotor’s contactable region. As with many previous models [9, 17, 35] our model places actomyosin within a negligible inertia, overdamped regime. As such we assume linear (viscous) drag. The resistive forces in the model will be referred to in terms of *α* and *β*, dimensionless constants representing viscous drag and the elastic returning force, respectively. The derivation of these constants and the values used to determine their ranges can be found in the methods and Table.I.

**TABLE I.** Parameter values taken from literature used to inform numerical simulations.

| Parameter | Physical Meaning | Reference Values |
| --- | --- | --- |
| $F_m$ | Maximum power stroke force of Myosin | $3 - 11.5pN$ [16, 31, 32, 72, 76] |
| $f_N$ | Uninhibited Attachment rate of Myosin | $3 - 55s^{-1}$ [77–79] |
| $\kappa$ | Elastic returning constant | $0.16 - 0.31pNnm^{-1}$ [9, 12] |
| $C_d$ | Drag Coefficient | $0.01\mu Ns^{-1} - 50\mu Ns^{-1}$ <sup>a</sup> [9, 17] |
| $V_m$ | Maximum Gliding Velocity of Unloaded Actin | $0.9 - 7\mu Mms^{-1}$ [30, 78] |
| – | Myosin Step Size | $5 - 17nm$ [68–73] |
| $l_r(physiological)$ | Physiological spacing between adjacent myosin heads | $42.9nm$ [36] |
| $l_t(physiological)$ | Physiological spacing between adjacent actin binding sites | $38nm$ [18] |

<sup>a</sup> The lower bound here is estimated from the hydrodynamic drag on micron scale particles, and the upper bound is taken from the required drag found by other models to elicit sawtooth oscillations.

Here we demonstrate that discontinuous motor-backbone contact interactions via force balance are sufficient to entrain generic, linear force-velocity motors. The choice of a linear decaying force-velocity relation as our driving force for our rotors later allows us to build a physical DCR experiment using electric motors (for more detail see methods).

### A. The DCR model recapitulates emergent cooperativity of actomyosin complexes

To investigate myosin coupling dynamics we adjust the model geometries to closely represent the actomyosin complex. The periodic gaps between myosin heads and actin binding sites were set to 43*nm* [36] and 38*nm* [18], respectively. *In vitro* experiments on actomyosin systems predominantly incorporate a resistive elastic component - the actin filament is bonded to a bead suspended in a laser trap for minimal experiments (Fig. 2 a) [9, 12], whilst sarcomeres act against the compliance of other sarcomeres and often against a flexible micro-manipulator needle when observing SPOCs in myofibrils (Fig. 2 b) [8, 34]. We therefore added a linear elastic returning force alongside the viscous drag to represent these spring-like components [8, 9, 17, 34]. The active number of heads used was 100. Finally, as a myosin head tends to spend longer in non-power stroke states than carrying out a power stroke [37] we set the contactable region in the rotors arc ‘*c*’ to 0.22 (see Methods). This effects how long the rotor has to travel before it returns to the contactable region, implemented by increasing the size of the rotor radius while maintaining the same length contactable region between tooth and rotor (Fig. 2 c). As the radius is increased we see that oscillations are more reliably generated (Fig. 2 c). A complete description of the models implementation as a numerical simulation is included in the Methods section and Eqs. (1-8).

In Fig. 2 d we show two examples of emergent behaviours of our system that closely resemble the specific cooperativity of the *in vitro* actomyosin results shown in (Fig. 2 a, b). These have been chosen from an exploration of varying *α* and *β* (*α* ∈ (0.2, 4) and *β* ∈ (0.1, 3)) and are qualitative matches chosen without fitting. It is a powerful demonstration of the importance of the mechanical regime the actomyosin geometry and architecture lie in, that both of these behaviours have been recreated with 100 rotors, when the first experimental result is produced with 17 myosin heads [12], and the second has many thousands. This has much to do with the form of the force balance equation (Eq. 1) where the forcing term scales with the number of myosin heads, *N*. Therefore as long as *α* and *β* are also scaled with *N* the behaviour will remain broadly the same (see supporting information). In the first example, (Fig. 2 d, i-iv) *α* = 1 and *β* = 3, chosen as the displacement is similar to a minimal actomyosin experiment (Fig. 2 a). The displacement profile is formed of many highly irregular saw tooth peaks. The rising segments of the oscillations are rough or stepped and the peak amplitude has a high variance (Fig. 2 d, i). The kymograph of the rotor positions shows a disorganised pattern moving into a roughly organised travelling wave (Fig. 2 d, ii). Taking a scatter plot of the number of rotors in contact with teeth for a given returning force shows the maximum number of rotors that can contact is 38% (Fig. 2 d, iii). This is a quality set by the geometry used. The scatter plot also shows a weakly increasing number of rotors contacting for an increasing returning force. We can see the highest returning force experienced by the system is 140*pN*. The moving probability density of the phases shows that the majority of motors are within *π* radians of their neighbour (Fig. 2 d, iv), and there are peaks of density around 0.2*π* radians. These peaks appear and disappear with no obvious accompanying features in the displacement profile. In the second example, (Fig. 2 d, v-viii) *α* = 2.4 and *β* = 0.5 the displacement profile is qualitatively closer to that of a spontaneously oscillating sarcomere (Fig. 2 b). Here the rising segment of the saw tooth peaks are smooth with a consistent gradient, and the amplitude of the oscillations is regular. The oscillations are slightly non-uniform, with variation in the depth of the fast falling segment of the sawtooth. The kymograph of the rotor positions (Fig. 2 d, vi) shows smooth metachronal travelling waves, with the rotors organised into sets of double peaks. These spontaneous travelling waves move in the opposite direction to the direction of the teeth. The scatter plot of the contacting rotors vs returning force is much tighter here (Fig. 2 d, vii), with a positive correlation between increasing force and % rotors in contact, and a maximum force of 135*pN* and 36% of motors in contact. There is also a more obvious grouping of points with few or no rotors in contact at higher forces (Fig. 2 d, vii, blue box). These points are related to the fast falling segment of the sawtooth when the system drops backward. The moving probability of the phases (Fig. 2 d, viii) shows a strong peak around *π/*2 radians and a weaker peak at 0 radians. The peaks disappear and the phases spread out very briefly at the fast falling segments of the sawtooth oscillations.

### B. Motor synchronisation and cooperativity are orchestrated by the actomyosin network’s architecture

There are three main spontaneous synchronisation and cooperativity patterns instigated by the actomyosin architecture. This is obtained by varying the motor-coupling strength via *α* and *β*. Multiple patterns can occur for a single parameter value set, such as one of the examples chosen for (Fig. 3a) where two patterns are present. The patterns have been named singlets (video 4), doublets, and triplets due to the presence of one, two or three wave maxima groupings (Fig. 3a). The groupings are dictated by how many times a rotor completes a full revolution before coming into contact with a tooth. For example, with a doublet a given rotor will complete one full rotation without contacting (Fig. 2d, vi), then contact a tooth on the second rotation. A comparison between the moving probability density of neighboring phase separations of the rotors with the locations of the synchronisation patterns on the displacement graph (Fig. 3a) shows for singlets there is a very strong phase grouping at 0.2 − 0.3*π* radians. Doublets show a weaker and wider phase grouping around 0.5*π* radians (the same as those shown in Fig. 2d, viii) and triplets show a very weak and heavily spread grouping centred on 0.7 −0.8*π* radians. All patterns have an additional peak around 0 radians. Fig. 3b shows a parameter sweep showing the different behaviours of the system for values of *α* and *β*. Behaviour A is very low amplitude jittering, where motors are uncoupled. Behaviour B is a combination of low amplitude peaks interspersed with well synchronised high amplitude intermittent singlet peaks. Behaviour C is low amplitude irregular oscillations (well demonstrated in Fig. 2d, i). Behaviour D is given by consistently taller, more organised peaks, such as those in Fig. 2d, v). It is important to note that though the peaks in behaviour D are more organised, they do generally demonstrate irregularity in period and amplitude such as the doublet/triplet example in Fig. 3a. Examples of strong synchronisation are only found within the intermittent strong peaks of behaviour B and the more organised peaks of behaviour D. Finally, Fig. 3c shows the instantaneous (light blue) and moving average (dark blue) velocity and number of rotors in contact (red) against the elastic returning force from the examples of each synchronisation pattern in Fig. 3a. The patterns reach a maximum rotors in contact of 30 − 35%, and a maximum force of 100 − 130*pN*. Though for these examples the doublets and triplets have greater maximum forces than singlets, this is not the rule. The average velocity of a singlet for a given returning force is twice that of a doublet and three times that of a triplet. The examples in Fig. 3 have the moving average velocity of 1200*nm/s*, 600*nm/s* and 400*nm/s* at 90*pN* for singlets, doublets and triplets, respectively. These figures show singlets have the smoothest instantaneous velocity, with triplets having the most variable instantaneous velocity.

**FIG. 3.**
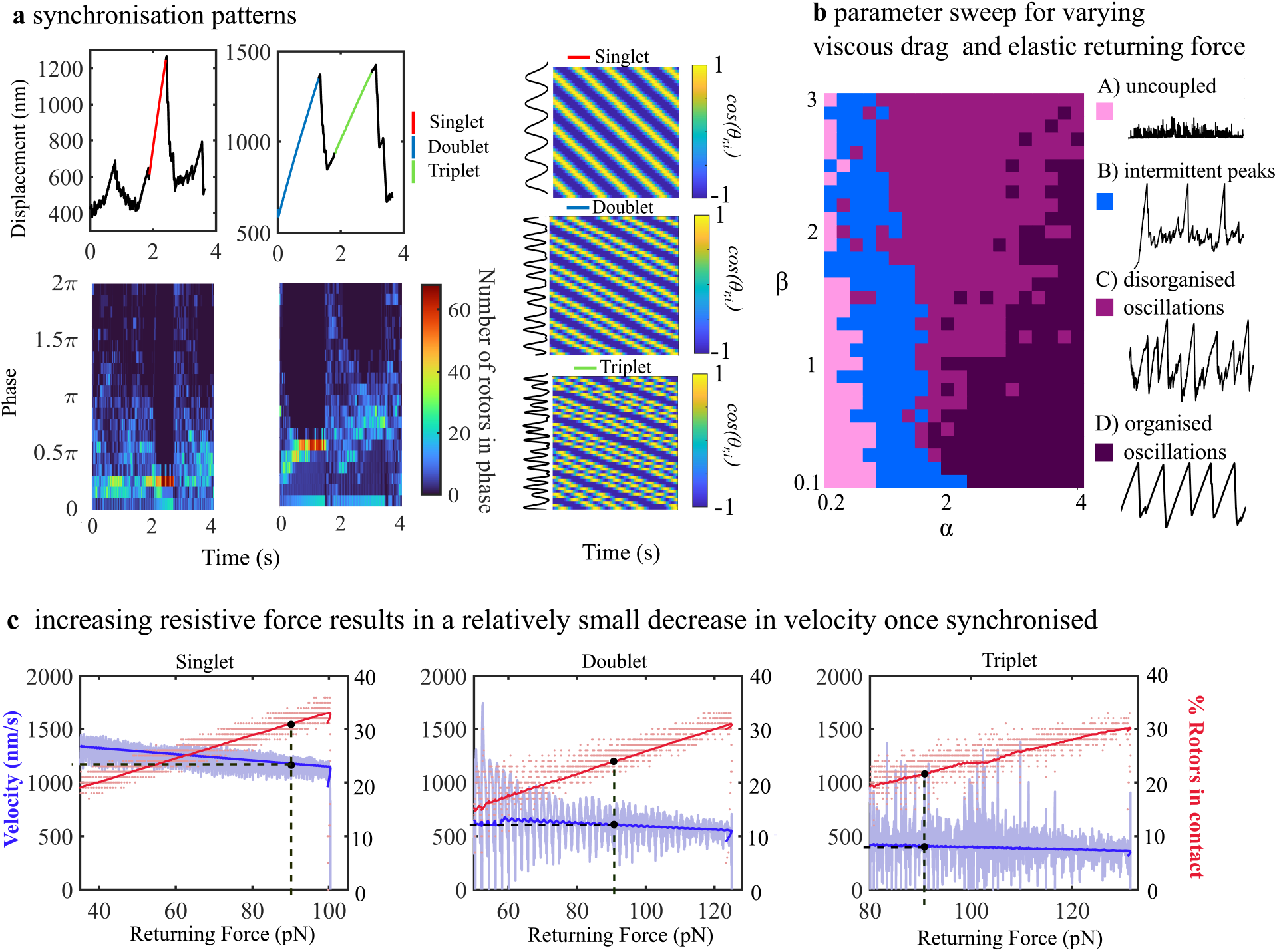
**a** Examples of the displacement profiles of three synchronisation patterns, with the accompanying rotor to neighbor phase difference probability distribution shown below and kymographs of the metachronal waves shown to the side. Singlets (shown in red) appear more reliably at lower viscous drag coefficients, and are often seen as smooth well organised spikes intermittently appearing in an otherwise disorganised signal. Doublets (blue) and triplets (green) appear more consistently when the viscous drag is higher. **b** A parameter sweep showing the locations of four key behaviours in the parameter space of varying viscous drag (*α*) and elastic returning force (*β*). **c** Velocity of the filament and percentage rotors in contact against elastic returning force for the three synchronisation patterns. The moving average velocity is shown in dark blue with the instantaneous velocity shown behind in light blue. Moving average of rotors in contact is shown in dark red, with the instantaneous number of rotors shown as a scatter in light red behind. The values at 90pN are highlighted for discussion in the text.

### C. Physical experiment of discontinuously coupled rotor network

A novel physical DCR experiment was constructed to study the emergent synchronization behaviour of motors through the discontinuous mechanical coupling that is instigated by the actomyosin architecture (Fig. 1e). Though we ensured the geometry was incommensurate, the weakness of the contact coupling (due to the semi-coupled, discontinuous nature of the system), in combination with a realistic limit to the number of motors we were able to use in the experiment meant we could not use the same physiological ratio of spacing between actin binding sites and myosin heads. Instead we adjusted to a geometry that allowed coupling with fewer motors (see Methods). The experiment starts with all rotors set to *θ* = 0, and as a consequence a short transient is present. Within the transient the displacement increases in a stepping fashion as the rotors contact and then disengage together (Fig. 4a, blue inset). Not all motors contact in the same position on their circular path due to the incommensurate nature of the rotors and teeth. This lets the rotor orientations spread out, settling to a phase difference of approximately 3*π/*2 between neighboring rotors (Fig. 4c). This results in a much smoother displacement increase (Fig. 4a, red inset). The outcome of a consistent phase difference between rotors is the spontaneous emer-gence of metachronal travelling waves that move in the same direction as the teeth (Fig. 4b, video 1,2). The simulation built to replicate the experiment expresses all of the same key attributes, but the transient is much shorter (Fig. 4d). The resulting travelling waves are qualitatively more ordered (Fig. 4e) with a more tightly packed phase probability density (Fig. 4f, video 3). This is likely attributable to the occurrence of noise and small perturbations due to the vibrations or imperfections present in the experiment. The parameters used in the experiment and how the simulation was adjusted to match them is detailed in the Methods section. The DCR experiment thus shows that a high level of cooperativity and synchronization between oscillators with self organised dynamics can be achieved via the mechanical organization that is imposed by the actomyosin architecture, without requiring intricate attachment-detachment reaction kinetics of biological motors and despite the time varying nature of the coupling.

**FIG. 4.**
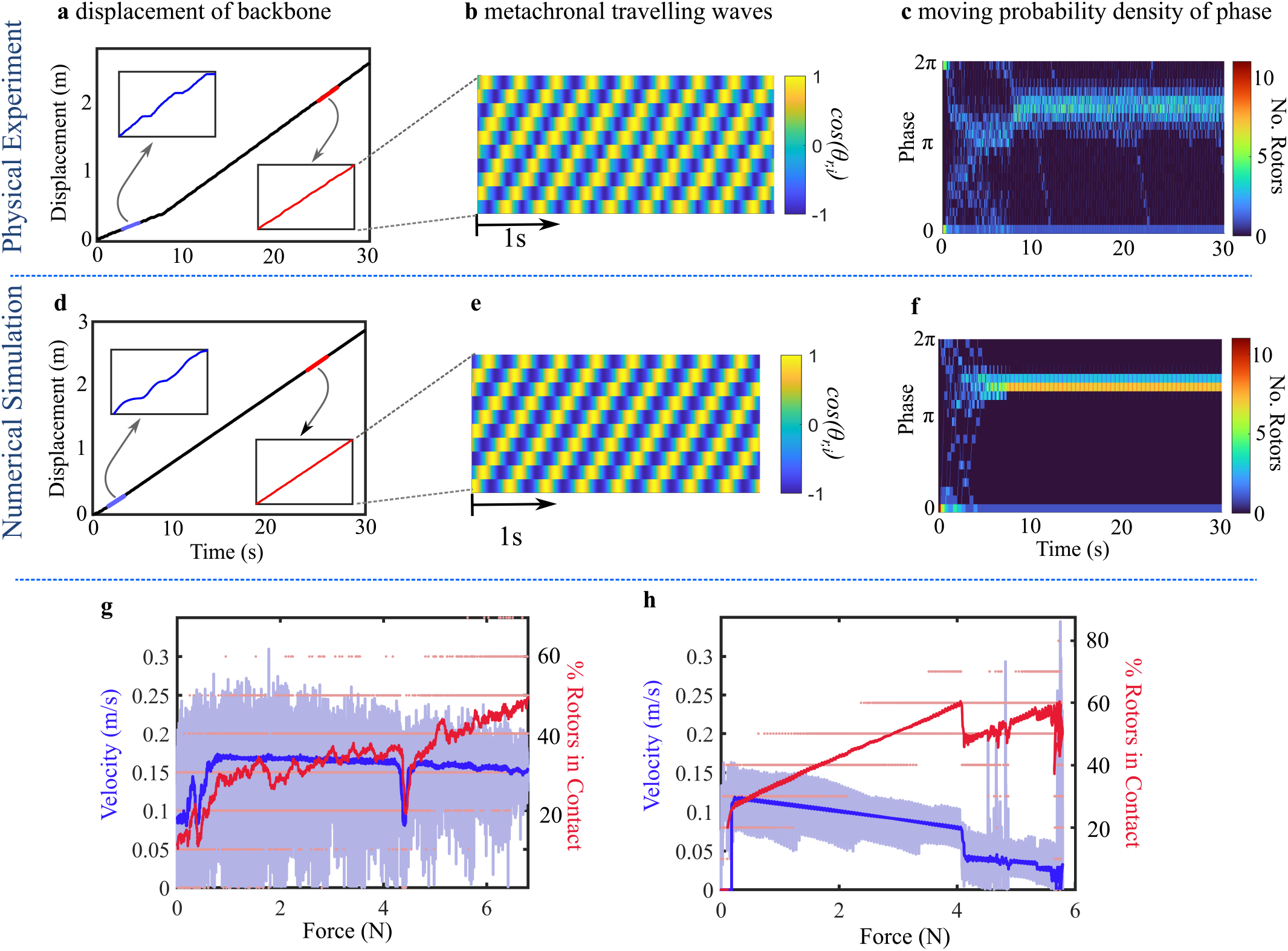
Comparison between kymograph, displacement profile and phase distribution for experimental (a-c) and simulated (d-f) systems. **a**,**d** The displacement smooths out after short transient, made obvious by the change in average gradient and reduction in stepping clearly seen in the displacement profile (blue inset to red inset). **b**,**e** A kymograph of the cosine of the rotors’ angle against vertical after the transient shows the development of travelling waves. **c**,**f** The phase distribution tightens into a band around approximately 3*π/*2 after the transient. The transient of the simulation is much shorter and the phase distribution is very tight in comparison to the physical experiment. This is likely because the simulation is a perfect system, whereas the experiment will have some compliance and imperfections in the construction and slightly varied frequencies and torques in the motors. g, h Velocity and rotors in contact vs returning force for the experimental (**g**) and simulated (**h**) models. Velocity shown in light blue, rotors in contact in light red. Moving averages are shown in a darker blue and red. Qualitative behaviour is shown to be similar (velocity reduces with force, contacting rotors increase) however the specific behaviour is quite different. This is likely due at least in part to the simulated model having idealised perfectly back-drivable motors, allowing it to slip more easily.

In a second case, the system is run against the friction components and a spring. The rotors remain organised as the force is increased, and the number of contacting rotors steadily increases with the extension of the spring, from a minimum average of one to a maximum of five (Fig.4g). The velocity slowly but steadily decreases in conjunction with this. When a simulation was run in the same conditions the velocity of the system is less, and the contact more rapidly increases to the maximum of 60% as the antagonist force is increased. At a returning force of 4*N* the pattern of the synchronisation changes from singlet to doublet, and the velocity decreases further (Fig.4h). The causes of the discrepancies between the experimental and simulated velocity are at least in part due to the use of idealised rotors in the simulation. The idealised rotors are perfectly back-drivable and have a force velocity relation where the stall torque occurs at a zero velocity. The real electric motors in contrast are geared, making them difficult to back drive, and have a slightly more flattened force velocity relation, where the velocity suddenly drops to zero at the stall toque from a higher than zero value. This allows the electric motors to apply the same force as the idealised rotors at greater velocities.

## III. DISCUSSION

By stepping away from the traditional population-based, two-state models [5, 9, 17–19], and instead, providing a simplified mechanical coupled network model, we have demonstrated that the actomyosin’s intricate and non-linear spontaneous oscillations (on a timescale greater than the natural oscillation period of individual rotors, and a length-scale larger than an individual motor’s step-size) can be derived mechanically by discontinuously coupled Kuramoto style phase oscillators that are only periodically coupled by a common backbone (due to the specific contact interactions imposed by the actomyosin architecture). Though our formulation is similar to that of the two-state models at the momentum balance level (*ibid*), as required for total force balance of the molecular motor system, our binding dynamics are fundamentally different. Frequency of binding and unbinding here is dictated by the actomyosin geometry via the self organised time variable network. In a stalled configuration, for example, a rotor can be prevented from ever ‘unbinding’ to a tooth due to the necessity of forward motion of the backbone to allow continued rotation of the rotor. This follows directly from how the myosin heads were abstracted in our model, as a continuously cycling oscillator. We stress however that the current understanding of binding dynamics of biological myosin motors are markedly distinct, as the unbinding probability increases with the force they experience and the biding probability depends on the number of unbound motors (*ibid*). Despite the very distinct motor binding dynamics, the cooperativity observed here appears to display the diversity of emergent oscillations that are more closely related with experiments than two-state continuous models reported thus far [5, 9, 18, 19]. We therefore suggest that the potential for complex cooperative behaviour of actomyosin complexes is mainly governed by the architecture and geometry of the actomyosin system in a time variable network of oscillators, and we emphasise the veracity of using a network such as this in molecular motor study.

By showing displacement results that can be qualitatively very similar to prior experiments both on minimal actomyosin systems [9, 12] and whole sarcomeres [8, 34], we demonstrate the importance of the local mechanical environment and geometry on the dynamics of actomyosin complexes. By building our simulation with motors with a simple linearly decaying force velocity relation and using that to inform a physically constructed experiment we have shown that an electric motor can be a reasonable parallel to a molecular motor due to this similar force-velocity relation (operating within an actomyosin geometry). We have shown that synchronising behaviour can be reliably achieved for a large parameter range (Fig. 3b, behaviour D), and that the synchronised system is robust through a large range of forces, unless the maximum number of contacting motors is reached. A process has been demonstrated here where a small reduction in velocity of the actin strand due to increasing returning force gives more time for myosin head ‘binding’, more connections to occur between nodes. This allows an increasing overall output force when required, subsequently minimizing the velocity reduction, while also allowing minimising of required active units when the antagonist force is low. This suggests the emergence of a novel self-gearing and control mechanism (Fig. 2).

To mimic *in vitro* experiments, we use model parameters to the same geometry as observed in actomyosin complexes, 100 interacting motor heads, and have a hookian elastic returning force. We are able to show a variety of behaviours qualitatively similar to *in vitro* experiments on actomyosin systems by varying only two parameters, *α* and *β*. The primary behaviour we showed was the saw tooth oscillations of the system (Fig. 2d; i, v). These oscillations are a key response in cooperative non-processive actomyosin complexes. There are several proposed hypotheses as to why the system suddenly falls back. These hypotheses generally centre around force determined uncoupling, or a reduction in the binding potential due to thickening of the filament at high levels of contraction, causing filament separation [38, 39]. It is noteworthy that the latter does not account for the fall-back of the system in a minimal actomyosin experiment. In the DCR model, the saw tooth waveform is caused by a number of cooperating rotors always ‘bound’ and pushing the teeth, which allows a continuous displacement forward until a large ‘unbinding’ event occurs, causing a sudden spring-back of the system (video 4). In systems where synchronisation is not present this is an inconsistent process, where rotors cooperate more by happenstance than organisation. Events where all currently bound rotors release occur often, with large amplitude variations and within a small time frame. This produces high frequency, low amplitude irregular peaks such as in (Fig. 2d, i), closely mimicking the oscillations recorded by minimal actomyosin systems (Fig. 2a).

In systems where synchronisation occurs, such as Fig. 2d, v, the number of bound rotors increases until a maximum recruitment is reached, leading to a mass unbinding of rotors. The spring presents a relatively large returning force that, once untethered by the large number of motors, overwhelms and backdrives any few remaining motor that may otherwise be able to bind. This causes the sudden fall back of the system. Though not necessarily uniform in shape, peaks generated by synchronized motors tend to be much closer in height, and more consistently reach the maximum number of contacting rotors allowed by the actomyosin geometry (Fig. 2d, vii). These peaks seem to more closely mimic the displacement profile of a full sarcomere (Fig. 2b), though they do not consistently capture the regularity of SPOCs. We suggest this lack of consistent regularity is a result of the different structure of a sarcomere.

We therefore suggest that for a sufficient number of motors the oscillatory characteristics are heavily defined by the geometrical arrangement of the actomyosin complex, which determines the maximum recruitment of motors. This is however modulated by the motor-coupling strength given by the opposing forces, which determine whether the system is able to synchronise, provided a motor force-velocity response. This aligns with hypotheses that the varying oscillatory characteristics of actomyosin complexes are generally determined by the mechanical properties associated with their location in the body and structure [9, 40].

The framework presented here allows the investigation of internal oscillations driven by individual motors, as displayed in Fig. 2d, vi. It can be seen the simulated actomyosin specific DCR self-organises into spontaneous waves travelling internally along the complex. Interestingly, the travelling wave is moving in the opposite direction to the physical experiment in Fig. 4b. This is because the travelling wave is formed from the tooth-rotor interference fit, and for physiological actomyosin the periodic gap between myosin heads is greater than the gap between the binding sites, while the opposite is true for the physical experiment (Fig. 4b+e). The three synchronisation patterns we have shown (Fig. 3) all demonstrate a relatively small reduction in velocity of the backbone after a concurrent large increase in returning force. This could be taken as a form of control, where increasing forces reduces the velocity, allowing more rotors to engage and ultimately minimises the reduction in velocity. A close examination of Fig. 3c however demonstrates that this is a modified version of the motors’ force velocity relation. If we imagine continuing the force-velocity lines in Fig. 3c at the same gradient without an unbinding event, we would see them all stall around 500*pN*, the maximum theoretical force of 100 constantly bound rotors. Though this maximum force is the same, the maximum velocity of the backbone for all the synchronisation patterns is far below the maximum linear velocity of an unbound rotor (7000*nm/s*). This lower maximum velocity comes from only engaging rotors when necessary, and due to the smaller gradient, it does still reduce velocity loss compared to a system where all the rotors are engaged at all times. If we consider our macro-scale rotor experiment, it is hard to see the benefit of this mechanism over a rotary mechanical system with constant coupling of the motors and a speed reducing gear ratio, which could doubtlessly produce a higher maximum force, with a shallower force-velocity profile. When we consider the myosin molecule as the forcing unit however the benefit is more clear. Myosin only performs a power-stroke when bound to actin, and by the nature of its back and forth motion can only be periodically bound. If all myosin was bound at once there would be no way of creating a continuous motion and the sarcomere would stall. Self-regulating the number of bound molecules depending on the force the system must act against allows a continuous motion through a range of resistive force, despite the quite binary force production of the myosin heads, whilst also being an efficient use of energy. The mechanism described here is velocity dependant engagement and, as our model is deterministic, is a deterministic process. For any given synchronisation pattern we can predict the number of engaged rotors by the velocity. Despite this, velocity dependant engagement could still be a mechanism for varying myosin attachment rate for the more stochastic physiological system. If an antagonist force is placed on an actin filament of an actomyosin complex, it will cause the system to move more slowly. This means a binding site passing nearby to a free myosin spends longer within a ‘bind-able’ area and the myosin is more likely to bind, meaning a decrease in velocity leads to an increase in bound myosin heads. Thus, even when binding is a stochastic process, over a large number of myosin heads, we could potentially observe a similar relationship.

Fig. 3b shows synchronisation only reliably occurs (behaviour D) within the parameter space when the viscous drag is high enough in relation to the springlike returning force. The importance of viscous forces in synchronised and cooperative behaviour in this model is an interesting but not unexpected quality. The required values are greater than models for viscous drag within the sarcomere would suggest [41], however this aligns with the result found by Placais *et al* [9], who found the viscous friction coefficient required to obtain the expected coupling dynamics within their model was 2 to 3 orders of magnitude higher than predicted, whilst Julicher and Prost [17] used a damping coefficient assuming a viscosity 10^2^ to 10^3^ greater than water. The greater than expected restrictive force on filament movement is no longer generally thought to be caused just by the drag of negatively strained motors [28]. Instead it is attributed to protein friction in the filaments - the close proximity of proteins is thought to constrain motor protein speed [42], as well as friction intrinsic to the moving components of the motor itself [43, 44].

The results from the experiment in Fig. 4 provide compelling evidence that a physically constructed geometry of incommensurately spaced passive and active components, even if not the same as actomyosin, can orchestrate synchronisation and cooperativity among motors, despite only short periods of motor coupling (via rotor-tooth contact at the backbone). (Fig. 4a) demonstrates that organisation of the rotors changes the displacement from (i) a stepping motion in a less ordered transient to (ii) a smoother and faster displacement when organised. The generation of spontaneous metachronal travelling waves (Fig. 4b) as the steady state of this system show the high level of rotor cooperativity that can be induced by the this form of architecture. Metachronal waves appear frequently across all branches of cell motility studies [45–49], as an effective way to produce a consistent force over a relatively large length scale, using many small individual force producing units. The occurrence of nonlinear and discontinuous coupling in the rotor experiments and excellent replicability of the results with the numerical simulation, validates the DCR model and gives credence to the complex emergent cooperativity for the specific actomyosin simulations.

Our experiments also demonstrate the velocity dependant engagement present in the simulations (Fig. **??**g). As this self control mechanism is the best example from this system of morphological computing it makes a particularly pertinent and appealing illustration of the systems significance to the area of soft and contemporary robotics [50, 51]. Morphological computing and embodied intelligence are emerging technologies of interest, with the objective of making artificial muscles and robots more adaptive, efficient and versatile [52, 53]. Creating adaptive self-organising force producing units and actuators capable of dealing with an unpredictable environment is a priority in furthering technologies [51]. We hope that cooperative motor systems such as the DCR model proposed here could improve methods for capturing real muscle-like actuation and behaviour using non-biological motors, as current technologies mimicking muscles or actomyosin complexes employ overly prescriptive ratcheting mechanisms [54, 55].

From a mathematical standpoint this network is also novel among examples of time variant networks for two reasons; Firstly due to the emergent way the coupling of nodes occurs. Not only do the oscillators synchronise, the nodes’ *connections* self-organise into periodicity through the accumulative mechanical interaction of the complex via the total internal momentum balance of the system. Synchronization of time-varying networks are highly dependant on the dynamical nature of the nodes’ connections [27], and rewiring or vanishing of such connections in prior models is generally prescribed by some time dependant function [56, 57], set using pre-existing data [58], the connections change probabilistically [59–61] or some combination of the three [62]. The actomyosin time-dependent network is however coupled with the phase oscillator synchronization, leading to connections appearing and disappearing in an unprescribed but self-organising manner. Coupling between two simultaneously contacting rotors through the stiff backbone is extremely strong, but Fig 2d vii shows synchronisation occurs for as little as 15% overall connectivity in the network. The second reason this system unique in the literature is the physical realisation of its nodes as Kuramoto style phase oscillators. In the DCR model each node is a rotor, each connection occurring through the physical pathway of rotor-backbone-rotor. Prior ‘reallife’ literature examples such as social networks [62], neural networks [58] or even epidemiological [63] networks lack this form of direct physical representation. We hope thus that our model will attract interest of mathematicians working on the synchronization of time-varying networks [26, 27, 59].

## IV. CONCLUSIONS

Here we demonstrate that non-biological motors (rotors) cooperate in a similar fashion to biological motors when working within the actomyosin geometry; without requiring specific biochemical reaction kinetics governing the (un)binding of motors to actin. The geometry and architecture of the actomyosin complex is what orchestrates the motors. As such, the geometrical embedding of generic motors is the only requirement for the emergence of the observed diversity of spontaneous oscillations. Remarkably, model predictions are able to mirror distinct actomyosin and sarcomeric *in-vitro* experiments [9, 12, 34, 35]. The emergence of coperativity is further demonstrated with a macroscopic physical replica using non-biological motors that can also produce metachronal waves. This indicates that (i) complicated biochemical reactions from biological motors may not be needed to recreate muscle-like behaviour, (ii) the architectural design of the actomyosin complex drives cooperativity and, as such, the actomyosin complex may not rely on motor-specific qualities to achieve its biological function. This may be a linked with the evolution of the skeletal actomyosin geometry itself, and have an great impact on better understanding the mechanics behind some myopathies [64–66], and perhaps aiding in future treatments. Finally, (iii) the results presented here opens new possibilities for the use of non-biological motors in bio-inspired architectural designs of artificial muscles that are able to mirror self-organisation of natural muscular systems.

## V. METHODS AND MATERIALS

### A. Discontinuously coupled DCR model for actomyosin complexes in time variable networks

In our abstraction of an actomyosin system myosin heads are modelled as a discrete series of Kuramoto style phase oscillators (Fig. 1 c), also denoted as rotors, that are coupled mechanically via discontinuous contact-contact interactions to the common backbone (Fig. 1).

A node on the actomyosin network represents a myosin head undergoing a continuous cyclical motion (Fig. 1 c) within the actomyosin complex [67]. Each node in the actomyosin network contains an oscillator, represented by a point a radius away rotating around a circle. This point shows where each myosin head is in its oscillation cycle. Each oscillator circles at its own preferred rate when uncoupled, i.e. when disconnected in the network but at a rate dictated by the backbone if coupled. This does not necessitate coupled rotors are moving at the same rate, bound rotors at different stages of their rotation will progress with the same linear velocity but different rotational velocities (Fig. 1 c). The intermittent coupling is dictated by the brief periods of mechanical interaction among ‘myosin heads’ and the ‘actin binding’ sites within the actomyosin structure. Actin binding sites are modelled as periodic ‘teeth’ connected to a rigid backbone which are constrained to one dimensional movement and experience a passive resistive internal frictional force. This is representative of viscous and protein friction within the actomyosin structure (Fig. 1 d). When a myosin head is bound to the actin, an active node is connected to the network, and thus is able to exert a sliding force to the common backbone, representing the actin filament. Mathematically, we suggest the actomyosin complex is a time-dependent network [26, 27] in which node connectivity undergoes spontaneous oscillations. This model allows both a mathematical simplification and a mechanical derivation of the interaction among motors, which is dictated dynamically by the geometrical and mechanical properties of the time-dependent actomyosin network. Altogether, this simplification bypasses the need of prescribing complicated biochemical reaction kinetics for the emergent dynamics of each oscillator, and also maintains our oscillators as separate entities. This allows us to consider the coupling dynamics of the time dependant network that the motors are embedded within. Phase shifts emerge dynamically via spontaneous oscillations of the actomyosin coupling, and thus its network. Finally, the abstraction that is invoked here simplifies the resulting dynamical system enough to allow numerical tractability of actomyosin synchronization, as to solve a very large system of coupled differential equations with stochastic multi-parametric biochemical reactions is prohibitively time consuming.

Our governing equation takes the form of a simple forced spring damper equation, with the novelty of the model found within the forcing term. The forcing term is given as the sum of forces of rotors in contact, where contact is encoded in a heavyside function. The motors are regulated by contact, and as such all emergent cooperation is a result of the semi-coupled nature of the intermittent contact forces. To simulate our rotors we use the general force-velocity equation shared between electric and molecular motors, given by *T*_*t*_ = *T*_*s*_(1 − *ω*_*r*_*/ω*_0_), where *ω*_0_ is the unloaded rotational velocity of a motor and *T*_*s*_ is the stall torque. It is important to note the stall torque *T*_*s*_ is the maximum force a motor can exert, and as such if *ω*_*r*_ is negative, *T*_*t*_ = *T*_*s*_. The component of the force parallel to the actin filament is found with (*T*_*t*_*/R*_*r*_) | cos(*θ*_*j*_) |, where *θ*_*j*_ is the angular position of rotor ‘*j*’ measured from the positive *y* axis and *R*_*r*_ is the radius of the rotor’s path. *R*_*r*_ and *F*_*m*_, myosin’s maximum powerstroke force, are the lengthscale and force by which we non-dimensionalise. We use 1*/f*_0_ = 2*π/ω*_0_ as our timescale. A force balance equation can thus be written;

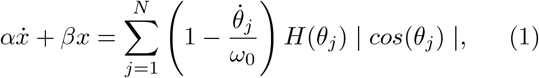

Where *N* is the total number of rotors, *α* and *β* are dimensionless constants for viscous drag on the filament and returning elastic force of the filament respectively, given by

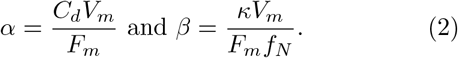

Where *V*_*m*_ = 2*πR*_*r*_*ω*_0_. Using the experimentally found parameter values for myosin systems shown in table.I, *α* ∈ (0, 116) and *β* ∈ (0.15, 13.15). The angular velocity of a rotor,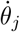, is given by

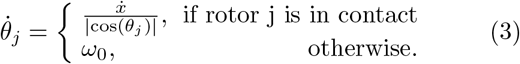

*H*(*θ*_*j*_) is a heavyside function indicating the contactable region’s projection of the rotors path;

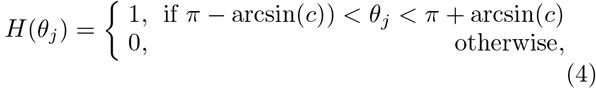

where ‘*c*’ is the dimensionless contactable region measured from the rotor’s centre (Fig. 1 b). The maximum step size of a myosin head is equivalent in this model then to twice a dimensional ‘*c*_*dim*_’, generally estimated between 8 − 17*nm* [68–73]. Contact of a rotor is determined when its *x* position falls within the contactable region and its separation from a tooth falls within a small set distance, ‘*d*’. A rotor’s *x* position is simply given as

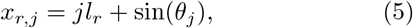

where *l*_*r*_ is the dimensionless length between rotor centre points. To find the location of teeth in the system first the number of teeth is found, 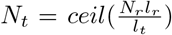, where *l*_*t*_ is the dimensionless length between teeth. The position of the teeth is then treated as a periodic,

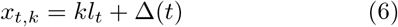

where Δ(*t*) = *X*(*t*)**mod** *l*_*t*_, therefore by comparing all rotor and tooth positions, contact can be established. In the situation where the teeth are springing back, all contacting rotors must have a negative rotational velocity and as such the rotors are assumed always exerting maximum torque, simplifying the original equation to

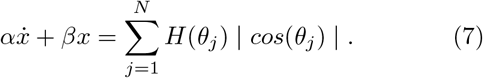

In order to solve equations (1 −7), they were broken into a matrix of coupled first order ODEs;

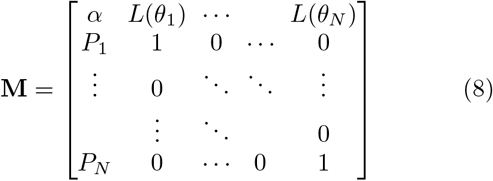

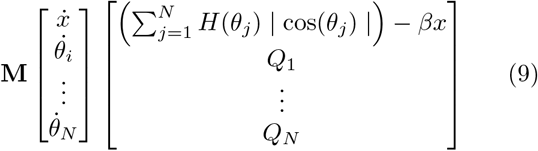

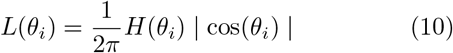

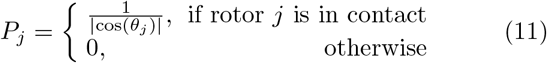

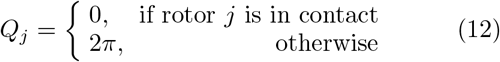

In the situation where the teeth are springing back, the rotors are always exerting maximum torque, simplifying **M** to

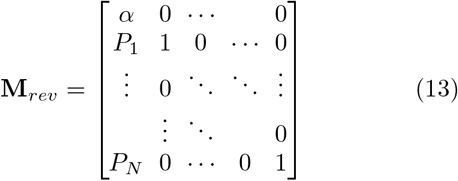

The equations were then solved using MATLAB’s ODE45 function. The MATLAB code is provided on GITHUB.

### B. physical experiment of DCR model

For our toybox model all custom components were designed in freeCAD and either 3D printed from standard PLA filament or laser cut from 5mm acrylic. Eleven 4Hz 6V geared motors were connected in parallel directly to a powerpack and were not limited by current draw. The motors are attached to rotors which spin and contact teeth, which are bonded to a belt with a serrated pin coated in cyanoacrylate glue. The belt drives a paddle which is submerged in a linearly viscous fluid (black treacle at 16 Celsius, estimated to have a viscosity of approximately 150*Pa*·*s* [74, 75] and the paddle had a surface area of approximately 0.6 ·10^−3^*m*^2^). To track the angle of each rotor and the total displacement of the belt the rotors and the driven shaft connected to the belt have targets. The targets have red dots which can be picked up by tracking software coded in MATLAB (Fig. 1 e). The geometry decisions taken to elicit synchronisation between the motors were taken to maximise the likelihood of synchronisation whilst minimising the required number of rotors. Building and maintaining a model large enough to house the number of motors needed to generate synchronisation and reliable travelling waves using the physiological myosin head separation to actin binding site separation ratio of 43 : 38 with a contactable region (*c*) of 0.2 was judged unreasonable for a proof of concept toybox model. Different ratios were tried with 11 motors, and an approximately 11 : 15 ratio with *c* = 0.9 was eventually shown to synchronise.

### C. Model parameters of the physical DCR experiment

The simulation was adjusted to reflect the parameters used for the experiment. The natural frequency, maximum torque, and rotor length are all known (176*rpm* [80], 0.02*Nm* [80], 0.015*m* respectively), therefore to calculate *α* we need to calculate the drag coefficient of the paddle and the paddle’s velocity in the fluid. The fluid was estimated to have a viscosity of approximately 150*Pa*·*s* [74, 75] and the paddle had a surface area of approximately 0.6·10^−3^*m*^2^, therefore *C*_*d*_ was found to have an approximate value of 0.9*N* ·*s*. The maximum velocity a rotor could cause the belt to move is 0.28*ms*^−1^. This is transferred at a 1:1 ratio to the average velocity of the paddle in the fluid giving *α* an approximate value of 0.2. This didn’t produce the expected smooth profile and travelling wave, so a small amount of static friction was added to the model in the form of a conditional constant, *γ*. Considering there was no returning force, and as such the displacement was always positive, *γ* was added as a constant resistive force that acted against the direction of travel of the filament. With *γ* set to 0.2, the expected pattern was formed.

### D. Model parameters of the actomyosin simulations

As there are a large number of free parameters in this model, as many were set by literature values as possible. The simplest to set are *l*_*r*_ and *l*_*t*_, which have dimensional values of approximately 43*nm* [36] and 38*nm* [18] respectively. The step-size here is taken to be 12*nm*, and as such *c*_*dim*_ is 6*nm*. The effect of decreasing the ratio of *c*_*dim*_ to the rotors radius *R*_*r*_ is to increase the time the rotor spends unable to contact a tooth, equivalent to a refractory period. Choosing *c* is therefore an important decision, with larger values favouring faster synchronisation and smaller values allowing oscillations due to a longer un-contactable period (Fig. 2 c). For all simulations in this paper the experimental value of 7*µms*^−1^ was used as the value for maximum unloaded stepping velocity of myosin given experimentally by Uyeda *et al* [30] for an experimentally measured unloaded actin sliding assay. This was taken to be the maximum tangential velocity of a rotor, allowing us to use

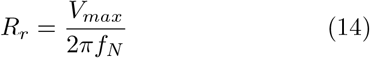

with a myosin attachment rate of 40*s*^−1^[77] in place of *f*_*N*_, to find the dimensional value *R*_*r*_ = 27.8*nm*. Taking the stepsize of a myosin head to be 12*nm* and when lengthscales are non-dimensionalised by 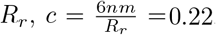. Using equations (2) and the literature values found within table. I we find *α* ∈ (0, 116) and *β* ∈ (0.15, 13.15), however, too high a value of either *α* or *β* prevents any meaningful dynamics so the more conservative ranges of *α* (0.2, 4) and *β* ∈ (0.1, 3) were considered. To maintain consistency, and demonstrate the impact of the mechanical environment on the behaviour of the actomyosin system, 100 rotors were used for every simulation shown in this paper. For results from other numbers of rotors see the supporting information.

## ACKNOWLEDGMENTS

This work was funded by the DTP EPSRC, and carried out using the computational facilities of the Advanced Computing Research Centre, University of Bristol - http://www.bristol.ac.uk/acrc/. The authors would also like to thank Mr Paul Fuchter for the many enlightening discussions and help during the research of this paper.

## Notes

### Competing Interest Statement

The authors have declared no competing interest.

### Summary of Updates

Article has been changed to emphasise physical experiment as a seperate section to the bulk of the main paper. An additional experiment was performed adding a springlike resistive force. Figure 4 was updated to reflect this additional experiment.

